# Low frequency and rare coding variation contributes to multiple sclerosis risk

**DOI:** 10.1101/286617

**Authors:** International Multiple Sclerosis Genetics Consortium, Mitja Mitrovic, Nikolaos Patsopoulos, Ashley Beecham, Theresa Dankowski, An Goris, Benedicte Dubois, Marie-Beatrice Dhooghe, Robin Lemmens, Philip Van Damme, Kate Fitzgerald, Helle Bach Sondergaard, Finn Sellebjerg, Per Soelberg Sorensen, Henrik Ullum, Lise Wegner Thoerner, Thomas Werge, Janna Saarela, Isabelle Cournu-Rebeix, Vincent Damotte, Bertrand Fontaine, Lena Guillot-Noel, Mark Lathrop, Sandra Vukusik, Pierre-Antoine Gourraud, Till Andlauer, Viola Pongratz, Dorothea Buck, Christiane Gasperi, Christiane Graetz, Antonios Bayas, Christoph Heesen, Tania Kumpfel, Ralf Linker, Friedemann Paul, Martin Stangel, Bjorn Tackenberg, Florian Then Bergh, Clemens Warnke, Heinz Wiendl, Brigitte Wildemann, Uwe Zettl, Ulf Ziemann, Hayrettin Tumani, Ralf Gold, Verena Grummel, Bernhard Hemmer, Benjamin Knier, Christina Lill, Efthimios Luessi, Efthimios Dardiotis, Cristina Agliardi, Nadia Barizzone, Elisabetta Mascia, Luisa Bernardinelli, Giancarlo Comi, Daniele Cusi, Federica Esposito, Laura Ferre, Cristoforo Comi, Daniela Galimberti, Maurizio Leone, Melissa Sorosina, Julia Y Mescheriakova, Rogier Hintzen, Cornelia Van Duijn, Steffan Bos, Kjell-Morten Myhr, Elisabeth Gulowsen Celius, Benedicte Lie, Anne Spurkland, Manuel Comabella, Xavier Montalban, Lars Alfredsson, Pernilla Stridh, Jan Hillert, Maja Jagodic, Fredrik Piehl, Ilijas Jelcic, Roland Martin, Mireia Sospedra, Maria Ban, Clive Hawkins, Pirro Hysi, Seema Kalra, Fredrik Karpe, Jyoti Khadake, Genevieve Lachance, Matthew Neville, Adam Santaniello, Stacy Caillier, Peter Calabresi, Bruce Cree, Anne Cross, Mary Davis, Jonathan Haines, Paul de Bakker, Silvia Delgado, Marieme Dembele, Keith Edwards, Hakon Hakonarson, Ioanna Konidari, Ellen Lathi, Clara Manrique, Margaret Pericak-Vance, Laura Piccio, Cathy Schaefer, Cristin McCabe, Howard Weiner, Thomas Olsson, Georgios Hadjigeorgiou, Bruce Taylor, Lotti Tajoori, Jac Charlesworth, David Booth, The Australia and New Zealand Genetics Consortium, The Wellcome Trust Case Control Consortium 2, Hanne Flinstad Harbo, Adrian Ivinson, Stephen Hauser, Alastair Compston, Graeme Stewart, Frauke Zipp, Lisa Barcellos, Sergio Baranzini, Filippo Martinelli Boneschi, Sandra D'Alfonso, Andreas Ziegler, Annette Oturai, Jacob McCauley, Stephen Sawcer, Jorge Oksenberg, Philip De Jager, Ingrid Kockum, David Hafler, Chris Cotsapas

## Abstract

Multiple sclerosis is a common, complex neurological disease, where almost 20% of risk heritability can be attributed to common genetic variants, including >230 identified by genome-wide association studies (Patsopoulos et al., 2017). Multiple strands of evidence suggest that the majority of the remaining heritability is also due to the additive effects of individual variants, rather than epistatic interactions between these variants, or mutations exclusive to individual families. Here, we show in 68,379 cases and controls that as much as 5% of this heritability is explained by low-frequency variation in gene coding sequence. We identify four novel genes driving MS risk independently of common variant signals, which highlight a key role for regulatory T cell homeostasis and regulation, IFNγ biology and NFκB signaling in MS pathogenesis. As low-frequency variants do not show substantial linkage disequilibrium with other variants, and as coding variants are more interpretable and experimentally tractable than non-coding variation, our discoveries constitute a rich resource for dissecting the pathobiology of MS.

## Main text

Multiple sclerosis (MS; MIM 126200) is an autoimmune disease of the central nervous system and a common cause of neurologic disability in young adults (Compston and Coles, 2008). It is most prevalent in individuals of northern European ancestry and – in line with other complex, common disorders – shows substantial heritability (Binder et al., 2016), with a sibling standardized incidence ratio of 7.1 (Westerlind et al., 2014). Over the last fifteen years, we have identified 233 independent, common variant associations mediating disease risk by genome-wide association studies (GWAS) of increasing sample size (Andlauer et al., 2016; Australia and New Zealand Multiple Sclerosis Genetics Consortium, 2009; Baranzini et al., 2009; de Jager et al., 2009; International Multiple Sclerosis Genetics Consortium, 2013; 2011; Jakkula et al., 2010; Martinelli-Boneschi et al., 2012; Nischwitz et al., 2010; Patsopoulos et al., 2011; 2017; Sanna et al., 2010; Wellcome Trust Case Control Consortium, 2007). Cumulatively, these effects – including 32 mapping to classical human leukocyte antigen (HLA) alleles and other variation in the major histocompatibility (MHC) locus (Moutsianas et al., 2015; Patsopoulos et al., 2013; 2017) – account for 7.5% of *h*^*2*^*g*, the heritability attributable to additive genetic effects captured by genotyping arrays, with a total of 19.2% of *h2g* attributable to all common variants in the autosomal genome (Patsopoulos et al., 2017). MS is thus a prototypical complex disease with a substantial portion of heritability determined by hundreds of common genetic variants, each of which explain only a small fraction of risk (Sawcer et al., 2014).

As with other common, complex diseases where large GWAS have been conducted, we find that although common variants (minor allele frequency, MAF > 5%) account for the bulk of trait heritability, they cannot account for its entirety. Identifying the source of this unexplained heritability has thus become a major challenge (Manolio et al., 2009). Two hypotheses are frequently advanced: that some common variants show epistatic (i.e. non-additive) interactions, so that they contribute more risk in combination than each does alone; and that a portion of risk is due to rare variants that cannot be imputed via linkage disequilibrium to common variants present on genotyping arrays, and are therefore invisible to heritability calculations based on such arrays. The only evidence we have found for epistatic interactions between common MS risk variants is being between two HLA haplotype families in the MHC locus (Moutsianas et al., 2015). This lack of epistatic interactions is consistent with other common, complex diseases, both of the immune system and beyond (Altshuler et al., 2008). We have also found no evidence that mutations in individual families drive disease risk in genome-wide linkage analyses of 730 MS families with multiple affected members (Sawcer et al., 2005). These results indicate that neither epistasis between known risk variants nor mutations in a limited number of loci are major sources of MS risk. They do not, however, preclude a role for variants present in the population at low frequencies, which cannot be imputed but are likely to individually contribute moderate risk.

Here, we report our assessment of the contribution of low-frequency variation in gene coding regions to MS risk. We conducted a meta-analysis of 144,209 low-frequency coding variants across all autosomal exons, concentrating on non-synonymous variants, which are more likely to have a phenotypic effect. We analyzed a total of 32,367 MS cases and 36,012 controls drawn from centers across Australia, ten European countries and multiple US states, which we genotyped either on the Illumina HumanExome Beadchip (exome chip) or on a custom array (the MS Chip) incorporating the exome chip content (Patsopoulos et al., 2017), and which satisfied our stringent quality control filters (Figure S1 and Tables S1 and S2). The exome array is a cost-efficient alternative to exome sequencing, capturing approximately 88% of low frequency and rare coding variants present in 33,370 non-Finnish Europeans included in the Exome Aggregation Consortium (minor allele frequencies between 0.0001 and 0.05; Figure S2), and <5% of the extremely rare alleles present at even lower frequencies. Our study was well powered, with 80% power to detect modest effects at low-frequency (odds ratio OR = 1.15 at MAF = 5%) and rare variants (OR = 1.5 at MAF = 0.5%) at a significance threshold of *p < 3.5 ×10*^*-7*^ (Bonferroni correction for the total number of variants genotyped).

We first assessed the contribution of individual variants to MS risk by conducting a meta-analysis of association statistics across 14 country-level strata (Figure 1). We used linear mixed models to correct for population structure in 13 of these strata, estimated from the 16,066 common, synonymous coding variants present on the exome chip (i.e. variants with minor allele frequency MAF > 5% in our samples). We included population structure-corrected summary statistics for the remaining cohort (from Germany), which has been previously described (Dankowski et al., 2015). As expected, we saw a strong correlation between effect size and variant frequency, with rarer alleles exerting larger effects (Figure S3). We found significant association between MS risk and seven low-frequency coding variants in six genes outside the extended MHC locus on chromosome 6 (Table 1 and Figure S4). Two of these variants (*TYK2* c.3310C>G, p.Pro1104Ala, overall MAF 4.1% in our samples; and *GALC* p.Asp84Asp, overall MAF 3.9%), are in regions identified by our latest MS GWAS, and show linkage disequilibrium with the common variant associations we have previously reported (International Multiple Sclerosis Genetics Consortium, 2011). The remaining variants are novel and are in neither linkage disequilibrium or physical proximity to common variant association signals.

**Table 1.**
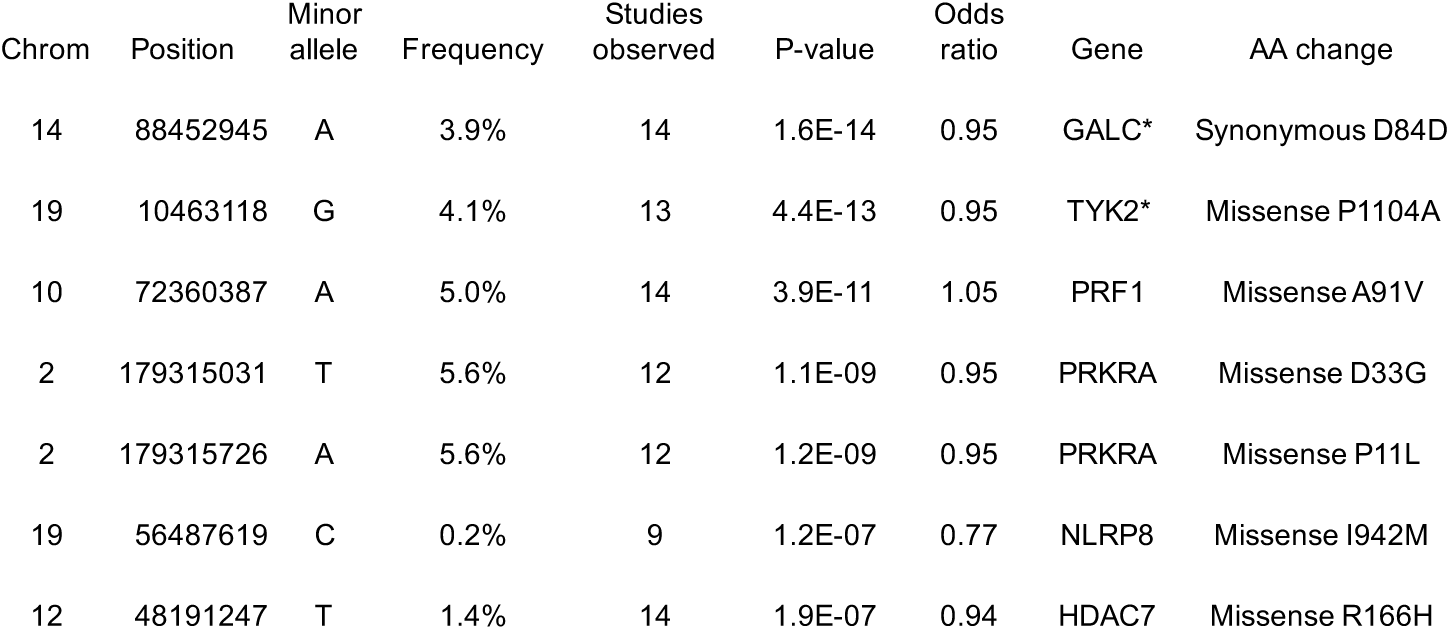
coding variants associated to multiple sclerosis risk. We analyzed 144,209 low-frequency non-synonymous coding variants across all autosomal exons in 32,367 MS cases and 36,012 controls drawn from centers across Australia, ten European countries and multiple US states. Genome positions are relative to hg19. The two variants in *PRKRA* are in linkage disequilibrium (R^2^ = 1, D = 1 in HapMap 3 European samples). * These variants lie in common variant risk loci found in our previous GWAS (Patsopoulos et al., 2017).

| Chrom | Position | Minor allele | Frequency | Studies observed | P-value | Odds ratio | Gene | AA change |
| --- | --- | --- | --- | --- | --- | --- | --- | --- |
| 14 | 88452945 | A | 3.9% | 14 | 1.6E-14 | 0.95 | GALC* | Synonymous D84D |
| 19 | 10463118 | G | 4.1% | 13 | 4.4E-13 | 0.95 | TYK2* | Missense P1104A |
| 10 | 72360387 | A | 5.0% | 14 | 3.9E-11 | 1.05 | PRF1 | Missense A91V |
| 2 | 179315031 | T | 5.6% | 12 | 1.1E-09 | 0.95 | PRKRA | Missense D33G |
| 2 | 179315726 | A | 5.6% | 12 | 1.2E-09 | 0.95 | PRKRA | Missense P11L |
| 19 | 56487619 | C | 0.2% | 9 | 1.2E-07 | 0.77 | NLRP8 | Missense I942M |
| 12 | 48191247 | T | 1.4% | 14 | 1.9E-07 | 0.94 | HDAC7 | Missense R166H |

**Figure 1.**
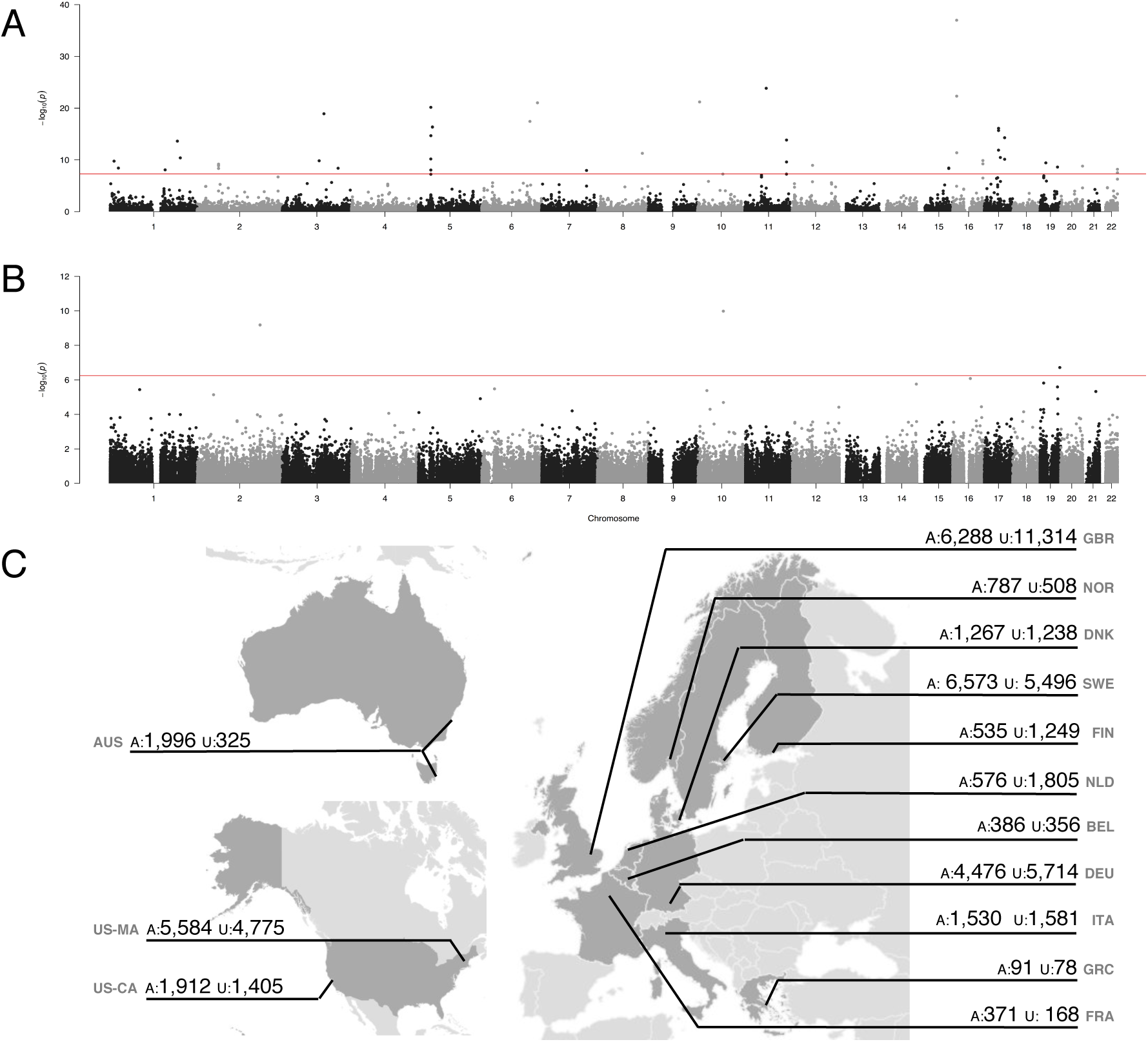
rare coding variants are associated to multiple sclerosis risk in a multi-cohort study. We analyzed 144,209 low-frequency non-synonymous coding variants across all autosomal exons in 32,367 MS cases and 36,012 controls drawn across the International Multiple Sclerosis Genetics Consortium centers. We find evidence for association with both common variants with combined MAF >5% (A); and with rare variants across the autosomes (B). We sourced samples from Australia, ten European countries, and the USA (C).

The newly discovered genes have clear immunological functions, confirming that MS pathogenesis is primarily driven by immune dysfunction. The associated polymorphisms show negligible linkage disequilibrium with other variants, so the genes harboring them are likely to be relevant to disease. *PRF1* encodes perforin, a key component of the granzyme-mediated cytotoxicity pathways used by several lymphocyte populations. In addition to cytotoxic lymphocytes and natural killer cells (House et al., 2015), perforin-dependent cytotoxicity is also seen in CD4^+^FOXP3^+^ regulatory T cells (Tregs), which show aberrant, T-helper-like IFNγ secretion in MS patients (Dominguez-Villar et al., 2011). The MS risk variant rs35947132 (p.Ala91Val) is associated with a decrease in target cell killing efficiency and increases in IFNγ secretion by NK cells (House et al., 2015), which aligns with the aberrant Treg phenotype observed in MS. This decreased cytotoxicity efficiency will prolong average cell-cell interactions with target cells, and such extended interactions are known to increase T cell receptor-mediated signaling and induce changes to T cell phenotypes, especially secretion of IFNγ and other cytokines (Constant et al., 1995). Similarly, *HDAC7* encodes the class II histone deacetylase 7, which potentiates the repressive effects of *FOXP3*, the master regulator governing naïve CD4^+^ T cell development into Tregs (Bettini et al., 2012; Li et al., 2007). *PRKRA* encodes protein kinase interferon-inducible double-stranded RNA-dependent activator; in response to double-stranded RNA due to virus infection, it heterodimerizes with protein kinase R to inhibit EIF2a-dependent translation, resulting in upregulation of NFκB signaling, interferon production and, eventually, apoptosis (Sadler and Williams, 2008). NFκB-mediated signaling is a core feature of MS pathogenesis, which we have shown to be altered by at least one MS-associated variant (Housley et al., 2015), and may be the relevant mechanism for this gene. Finally, *NLRP8* is an intracellular cytosolic receptor active in innate immune responses; the Ile942Met MS risk variant rs61734100 is detected only in individuals with European ancestry in ExAC.

Though we are able to identify individual low-frequency variants associated with MS risk, we recognize that we cannot detect all such variants at genome-wide significance. We thus sought to quantify the overall contribution of low-frequency coding variation to MS risk. In each of the thirteen strata that comprise our data, we estimated the proportion of heritability explained by common (MAF > 5%) and low-frequency (MAF < 5%) variants on the exome arrays (Yang et al., 2011). We included genotype-derived principal components to further control for population stratification. By meta-analyzing these estimates across the twelve strata where the restricted maximum likelihood model converged, we found that low-frequency variants explain 11.34% (95% confidence interval 11.33%-11.35%) of the observed difference between cases and controls (mean estimate 4.1% on the liability scale; Figure 2). We further partitioned the low-frequency variants into intermediate (5% > MAF > 1 %) and rare (MAF <1%), and found that the latter alone explain 9.0% (95% confidence interval 8.9%-9.1%) on the observed scale (mean estimate 3.2% on the liability scale; Figure 2). We note that six of the eight genome-wide significant variants presented in Table 1 are of intermediate frequency, and thus are not included in the rare category. Our results thus indicate that many more rare non-synonymous variants contribute to MS risk but are not individually detectable at genome-wide thresholds even in large studies like ours.

**Figure 2.**
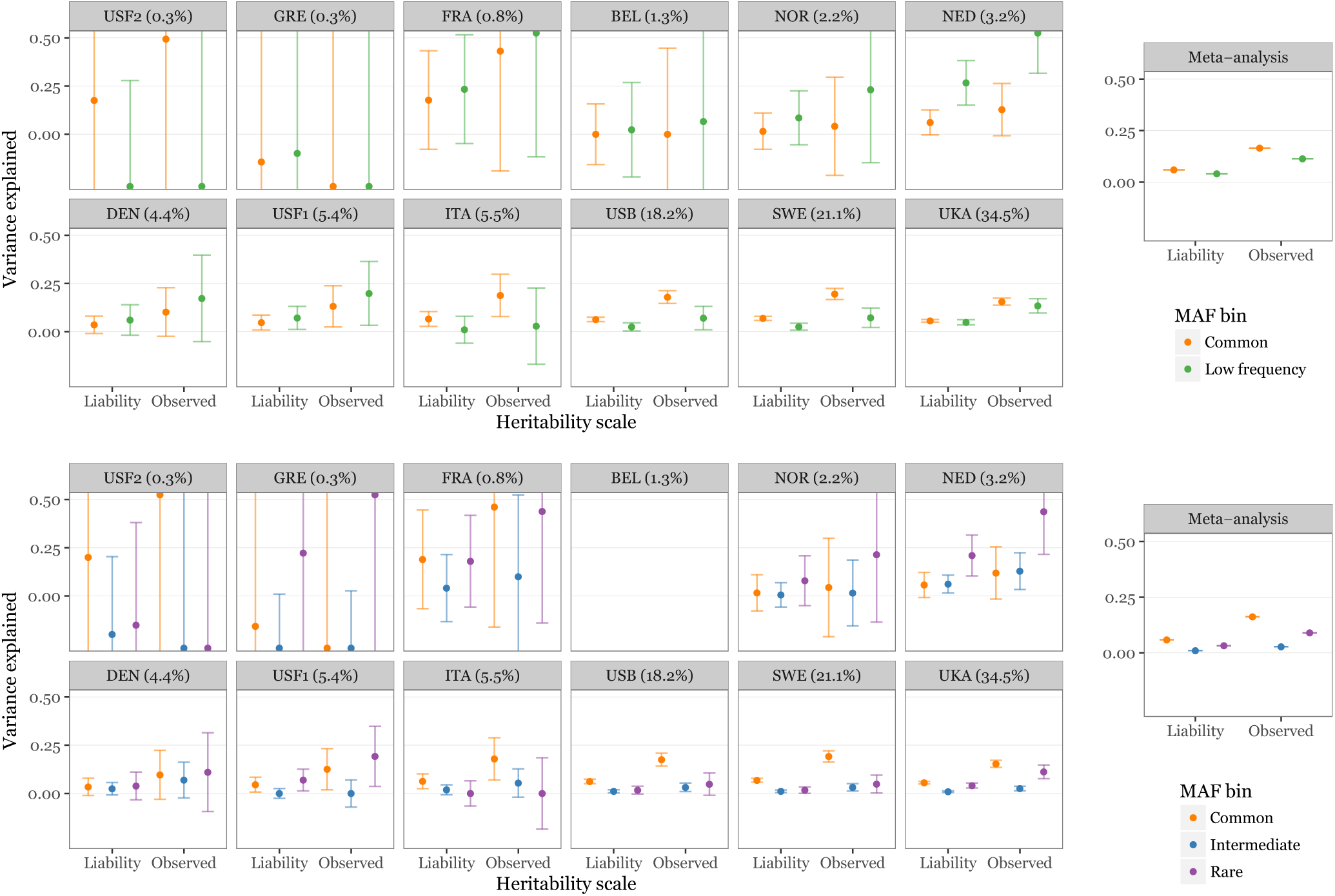
rare variants explain a substantial portion of multiple sclerosis heritability. We estimated the MS risk heritability explained by common variants (MAF > 5%) and low-frequency nonsynonymous coding variation (MAF < 5%) in each of thirteen cohorts genotyped on the exome chip, using GCTA (top panel). By meta-analyzing these estimates across cohorts, we found that low-frequency variants explain 11.34% of heritability on the observed scale, which corresponds to 4.1% on the liability scale (right top). After dividing the low-frequency variants into intermediate (5% > MAF > 1 %) and rare (MAF <1%; bottom panel), we found that the latter alone explain 9.0% heritability on the observed scale (3.2% on the liability scale; bottom right). Meta-analysis confidence intervals are small and visually occluded by the mean estimate plot characters. Cohorts (abbreviations as in Table S1) are ordered by sample size, with the percentage of the overall sample size shown in each subplot title. We could not obtain estimates for either model for our Finnish cohort (see Methods; not shown), or for the three-component model for our Belgian cohort (bottom panel, top row, fourth from left). Both cohorts are small, which may explain the failure to converge.

In this study, we show that low frequency coding variation explains a fraction of MS risk, which cannot be attributed to common variants across the genome. We capture most, but not all, low-frequency missense variants (Figure S2), suggesting our heritability estimates for low-frequency and rare variation are conservative. This broadly agrees with previous reports that such variants contribute to complex traits, including Alzheimer disease (Sims et al., 2017) and schizophrenia (Purcell et al., 2015), where heritability modeling similar to ours supports a role for rare variants. Studies of quantitative phenotypes shared by the entire population, such as height (Marouli et al., 2017), serum lipid levels (Liu et al., 2017) and blood cell traits (Chami et al., 2016; The CHARGE Consortium Hematology Working Group, 2016), have also reported novel associations to low-frequency coding variants outside the large number of known GWAS loci in each trait. However, a meta-analysis of different type 2 diabetes study designs found no associations outside common variant GWAS regions (Fuchsberger et al., 2016), though this may be due to the heterogeneity of sample ascertainment and study design. In aggregate, therefore, our results and these past studies demonstrate that rare coding variants contribute a fraction of common, complex trait heritability. These results also agree with both theoretical expectation and empirical observations that low-frequency coding variants are under natural selection, and are unlikely to increase in frequency in the population (Nelson et al., 2012; Schoech et al., 2017; Zeng et al., 2017). Thus, some portion of disease-associated variants, and hence the genes they influence, may not be detectable with conventional GWAS designs. Our discovery of multiple risk-associated genes that are central to IFNγ biology, Treg function and the NFκB signaling pathway in MS pathogenesis, and that do not reside in >200 known MS risk loci, supports this view.

## Acknowledgements

A full list of acknowledgements appears in the supplementary material.

## Methods

### Genotyping, quality control and stratum assignment

We assembled a total of 76,140 samples (36,219 cases, 38,629 controls and 1,292 samples with missing phenotype information) from across the International MS Genetics Consortium (IMSGC; Table S1). We genotyped these either on the Illumina HumanExome Beadchip (exome chip) or on a previously described custom array (Patsopoulos et al., 2017) including the exome chip content, both manufactured by Illumina Inc. We called genotypes both with Illumina’s default algorithm, gencall, and zCall, specifically developed to call low-frequency variants where all three groups of genotypes may not be observed (Goldstein et al., 2012).

An overview of our quality control process is shown in Figure S1; we used plink (Purcell et al., 2007) for all analyses unless otherwise noted. Briefly, we first excluded samples with low genotyping rate, extreme heterozygosity rate, inconsistent genotypic and recorded sex; we also removed closely related samples, keeping the relative with least missing data. Next, we removed population outliers by calculating genotype principal components using 16,066 common variants in linkage disequilibrium (r^2^ < 0.1) across the exome. We used EIGENSOFT 6 (Price et al., 2006) and FlashPCA (Abraham and Inouye, 2014) for cohorts with more than 10.000 individuals. We next removed variants with >3% gencall missing data rate for variants with minor allele frequency MAF >5%, or >1% zCall missing data rate for variants with MAF < 5%. We also removed variants out of Hardy-Weinberg equilibrium (p < 10^-5^). Next, we removed samples with high similarity in missing genotypes (“identity by missingness”) indicative of production artefact, and samples with missing phenotype information. Finally, we again removed any remaining population outliers using projection principal component analysis. We calculated 30 principal components for 1,092 individuals in 1,000 Genomes reference populations, again using the 16,066 common variants in linkage disequilibrium (r^2^ < 0.1) across the exome. We then projected the IMSGC samples into this space and excluded individuals more than six standard deviations from loading means as previously described (Price et al., 2006). We performed the projection and outlier detection and removal steps a total ten times to gradually remove more subtle population outliers.

We compiled cases and controls into strata for analysis as shown in Table S2. In total, we removed 17,951/76,140 (24%) samples either due to low data quality or as population outliers, leaving a final dataset of 27,891 cases and 30,298 controls in 13 strata (Figure S1 and Tables S1 and S2). Separately, we included summary statistics from 4,476 MS cases and 5,714 controls from Germany, genotyped on the exome chip as previously described (Dankowski et al., 2015), giving us a total of 32,367 MS cases and 36,012 controls for analysis.

### Exome chip coverage of ExAC variants

To assess how thoroughly the exome chip assesses low-frequency coding variation genome-wide, we compared it to the list of variants reported by the Exome Aggregation Consortium, ExAC (Lek et al., 2016), in their data release version 1. We filtered their summary table of all ExAC variants (available at ftp://ftp.broadinstitute.org/pub/ExAC_release/release1/manuscript_data/ExAC.r1.sites.vep.table.gz and last accessed 15 November 2017) for nonsynonymous coding variants passing their quality control, with at least one minor allele observed in non-Finnish European samples. We identified which of these variants are represented on the exome chip by comparing genomic coordinates (Figure S2).

### Univariate association analysis

We used mixed linear models for association analysis, as implemented in GCTA (Yang et al., 2011). In each of our 13 genotype-level strata, we calculated genetic relatedness matrices from 16,066 common, noncoding variants (overall MAF > 0.05) in linkage equilibrium (all pairwise r^2^ < 0.1) present on the exome chip, and with these calculated univariate association statistics for each autosomal variant present on the exome chip. To further control for population stratification, we also calculated genotypic principal components with the 16,066 common variants, and included these as covariates to the association analysis. We also included genotypic sex and chip type as covariates. We combined statistics across strata using inverse-variance-weighted meta-analysis, also as implemented in GCTA (Yang et al., 2011). As the bulk of exome chip variants are not common and do not show appreciable linkage disequilibrium, we controlled for multiple tests with a Bonferroni correction for the number of low-frequency variants, to give a genome-wide significance threshold of p < 3.58 *×* 10e^-7^ (0.05/139,764 variants with a combined MAF < 0.05 in controls and a heterogeneity index I^2^ < 50 in our meta-analysis).

### Heritability estimation

We used GCTA to calculate the heritability attributable to groups of variants in each of our 13 genotype-level strata (Yang et al., 2011). In each stratum, we ran two sets of models: a two-component model, estimating the heritability attributable to common and low-frequency (MAF ≤ 0.05) variants; and a three component model with rare (MAF ≤ 0.01), intermediate (0.01 < MAF ≤ 0.05), and common variants. In all strata, common variants are the set of 16,066 independent variants (overall MAF > 0.05) used for population stratification calculations in the univariate analysis above. We computed genetic relatedness matrices for each component of each model, then calculated narrow-sense heritability (*h*^*2*^) with 100 iterations of constrained restricted maximum likelihood (REML) fitting, assuming a disease prevalence of 0.001. We also included the principal components of population structure computed for the univariate analysis as covariates. As anticipated, several of the smaller cohorts presented fitting issues: no models converged for FIN; both three-component and two-component fits for UCSF2, and the three-component model for GRE would not converge under constraint and so were run without constraints; and the three-component model for BEL converged on two exactly equally likely solutions after 10,000 iterations. For the latter, we chose the most conservative estimates of variance explained. We combined these estimates with inverse variance-weighted meta-analysis.

## Supplementary information

**Figure S1.**
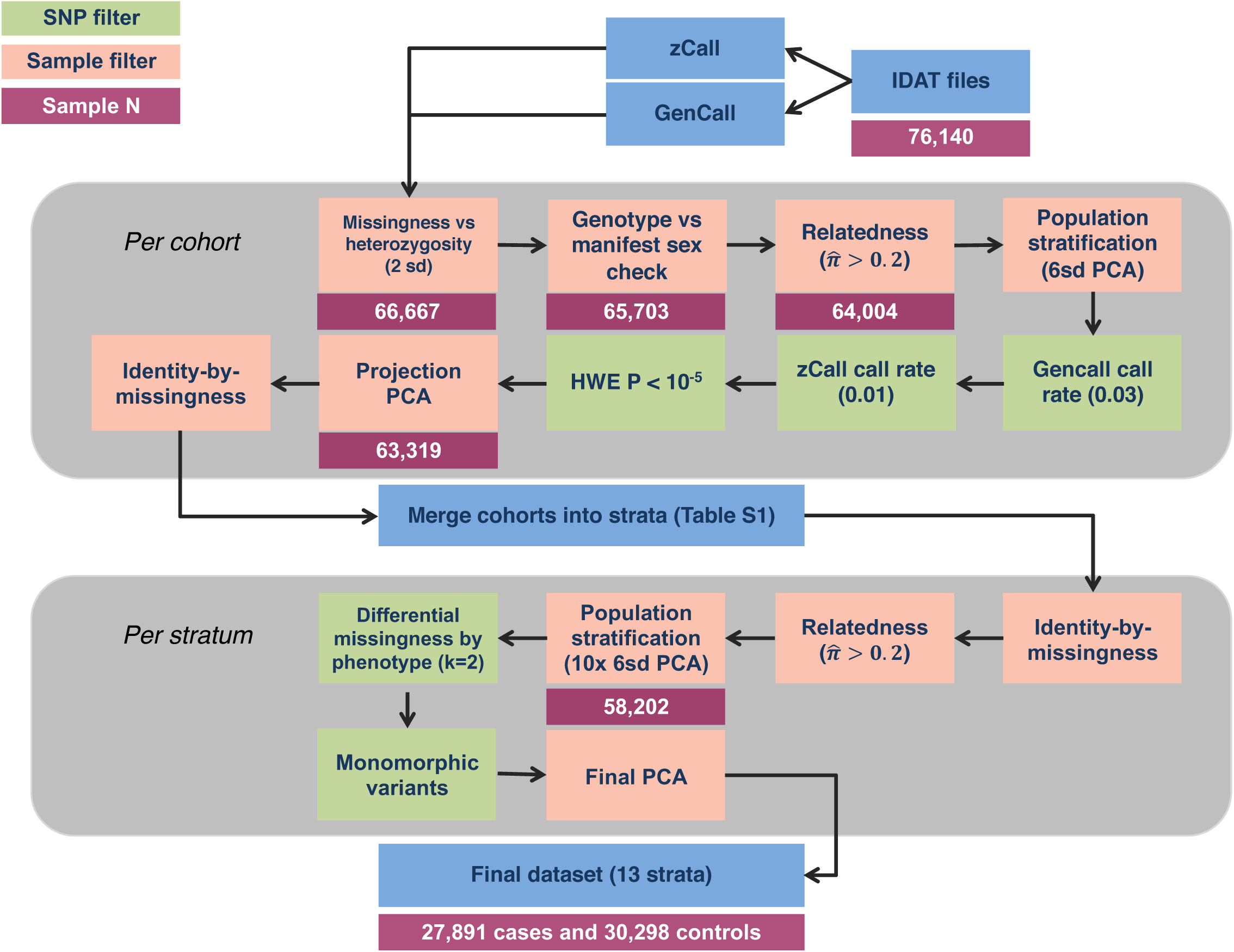
quality control pipeline overview. We assembled 46 cohorts of data (either entire country-level collections or groups of samples processed as a batch; Table S1). We called common variant genotypes with the standard algorithm provided by Illumina (GenCall), and low-frequency variants with zCall, an algorithm specifically developed to call these variants on the exome chip (Goldstein et al., 2012). We performed initial quality control on each cohort separately to account for variation between batches and cohorts (upper gray region), then merged cohorts into 13 country-level strata. To ensure that these strata were uniform we then performed stringent quality control on each stratum (lower gray region) to produce our final dataset.

**Figure S2.**
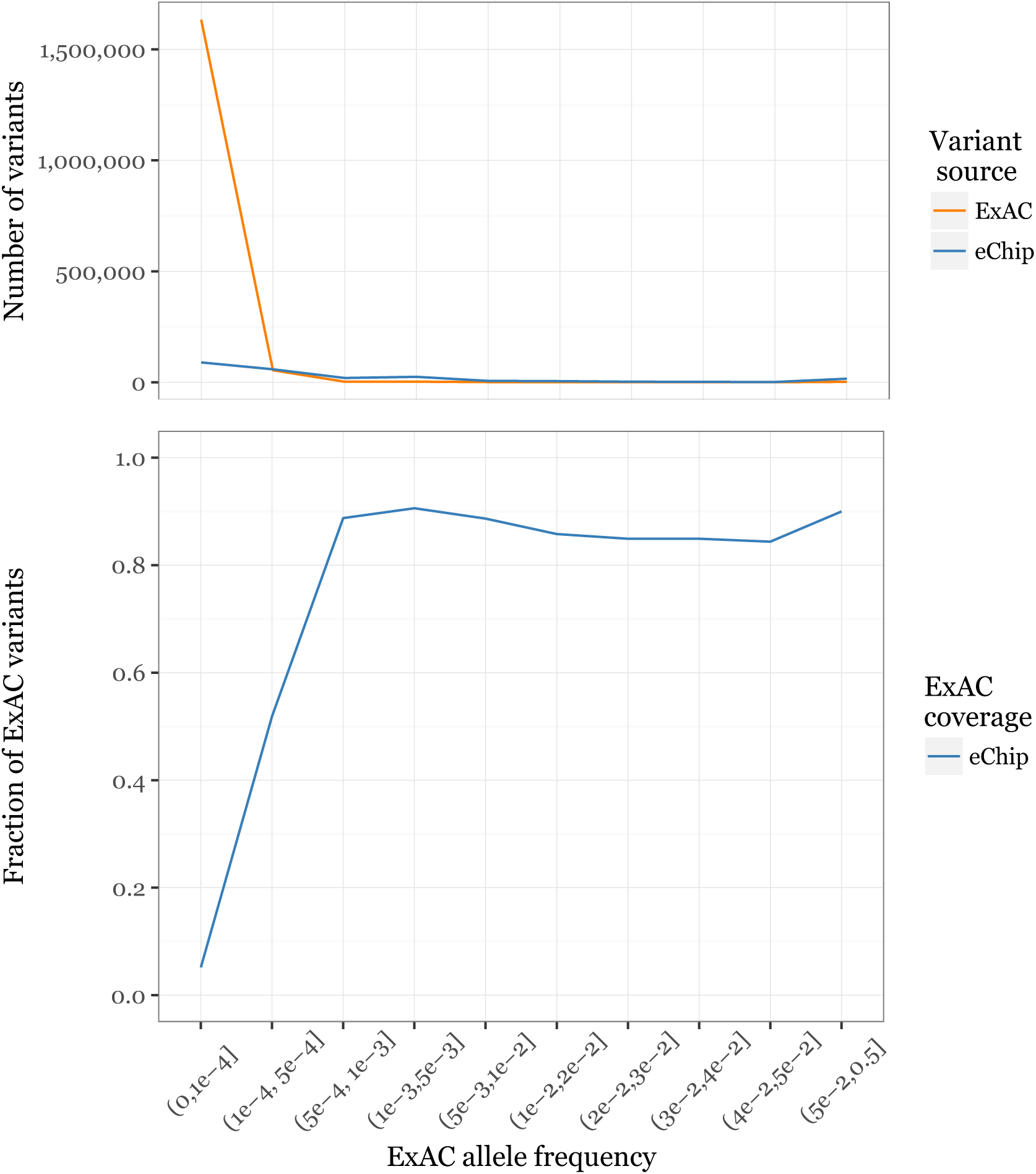
the exome chip captures a large fraction of ExAC (release version 1) low-frequency miss-sense variants. The exome chip captures the majority of variants present in ExAC (Lek et al., 2016) down to a minor allele frequency ∼ 0.0005, below which a large number of variants is observed (upper panel). Thus, the overall coverage at very rare alleles (5 × 10^-4^ > MAF > 1.5 × 10^-5^, corresponding to a single allele seen in 33,370 non-Finnish European individuals in ExAC) is low (lower panel).

**Figure S3.**
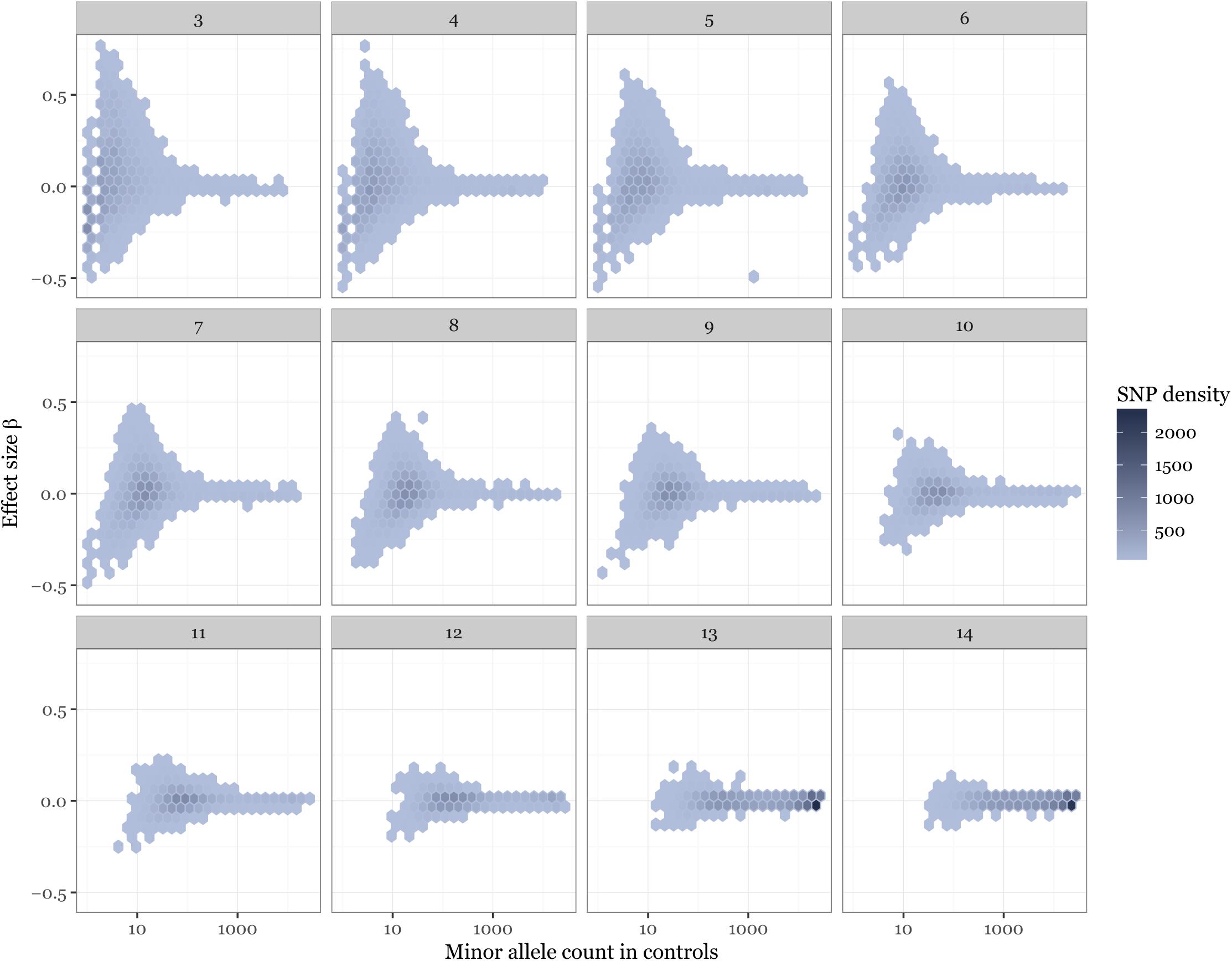
effect size correlates with minor allele frequency. We conducted a meta-analysis of 144,209 low-frequency coding variants across all autosomal exons, concentrating on non-synonymous variants which are more likely to have a phenotypic effect. We analyzed a total of 32,367 MS cases and 36,012 controls in thirteen strata. Here, we show that effect size (β or log odds ratio, y axis) correlates to allele frequency (number of minor alleles present in control samples, x axis). Because many low-frequency variants are not present in all cohorts, we stratify these data by number of cohorts in which a variant is polymorphic (subplots). Rarer variants have larger estimated effect sizes, and are present in fewer cohorts.

**Figure S4.**
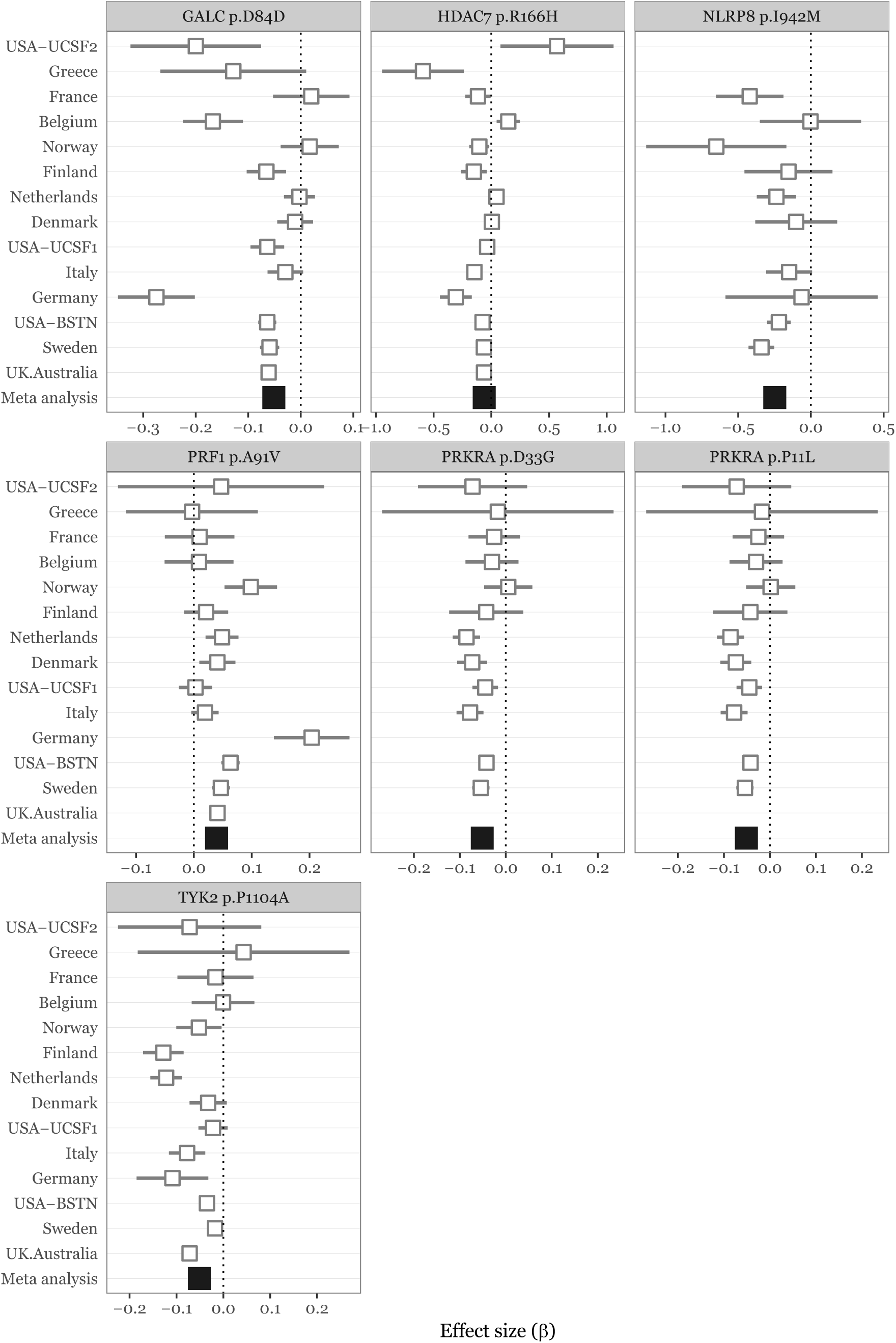
forest plots for genome-wide significant low-frequency variants. Seven variants in six genes are significant in our analysis (*p < 3.5* × *10*^*-7*^, Bonferroni correction for the total number of variants genotyped). Two of these (*TYK2* p.Pro1104Ala and *GALC* p.Asp84Asp), are in linkage disequilibrium with known GWAS hits. Studies are ordered by increasing sample size.

**Table S1.**
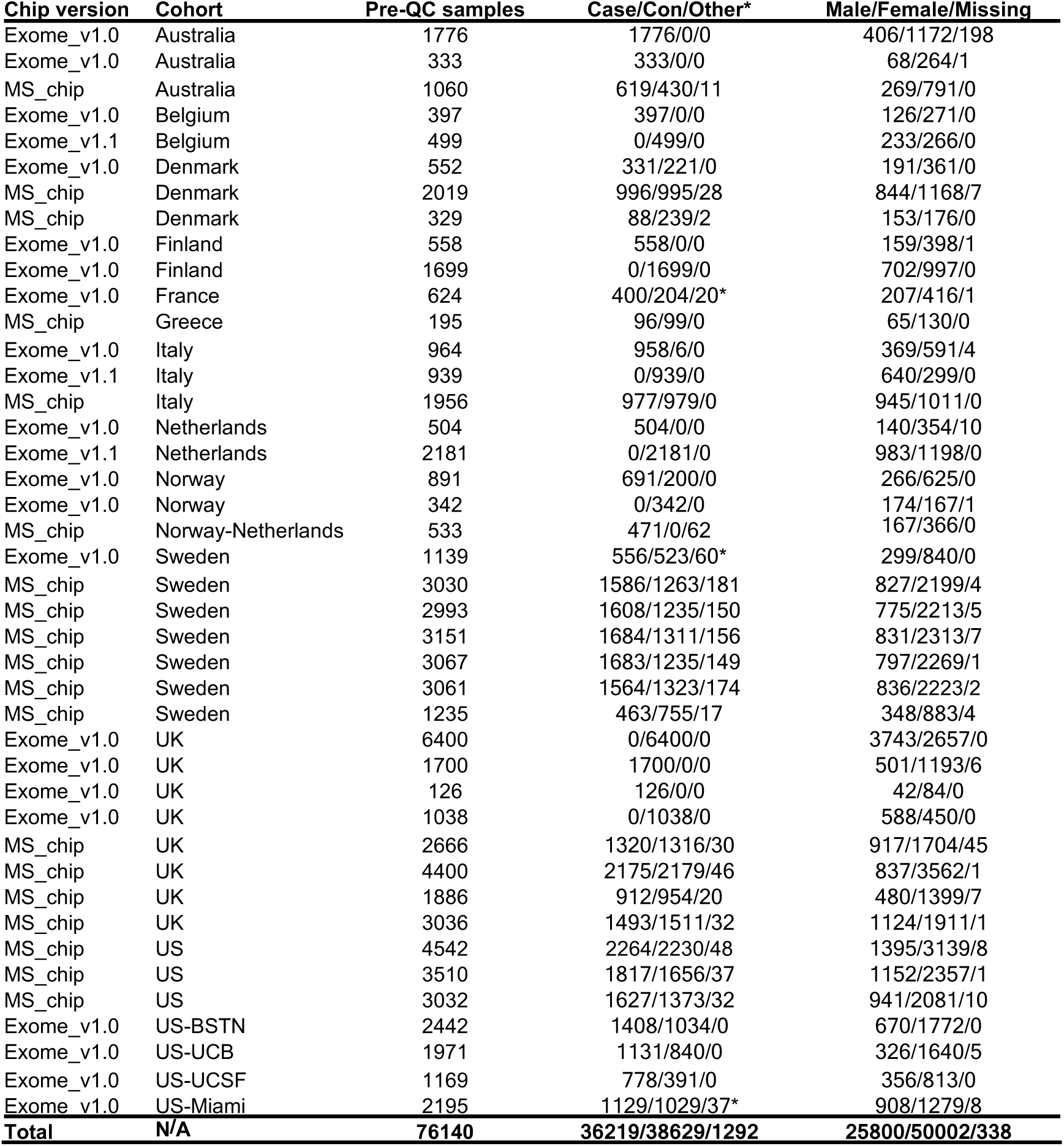
genotype-level samples included in our study. 58,189/76,140 (76.5%) of our samples passed quality control and could be assigned to one of thirteen strata. 9,473/17,951 (52.8%) of failed samples did not pass a heterogeneity versus missing data rate filter, suggesting either poor data quality or population stratification (detailed in Figure S1). We used two versions of Illumina’s HumanCore Exome array: the standard product (version 1.0; designated *Exome_v1.x* in the chip type column) and a customized version including ∼100,000 additional variants we specified (designated *MS_chip*), described elsewhere (Patsopoulos et al., 2017). Belgian control samples were genotyped at the Center for Inherited Disease Research (CIDR, Baltimore, MD, USA) on the Illumina 5M array (Illumina, San Diego, CA, USA) as part of the Stroke Genetics Network (SiGN).

**Table S2.**
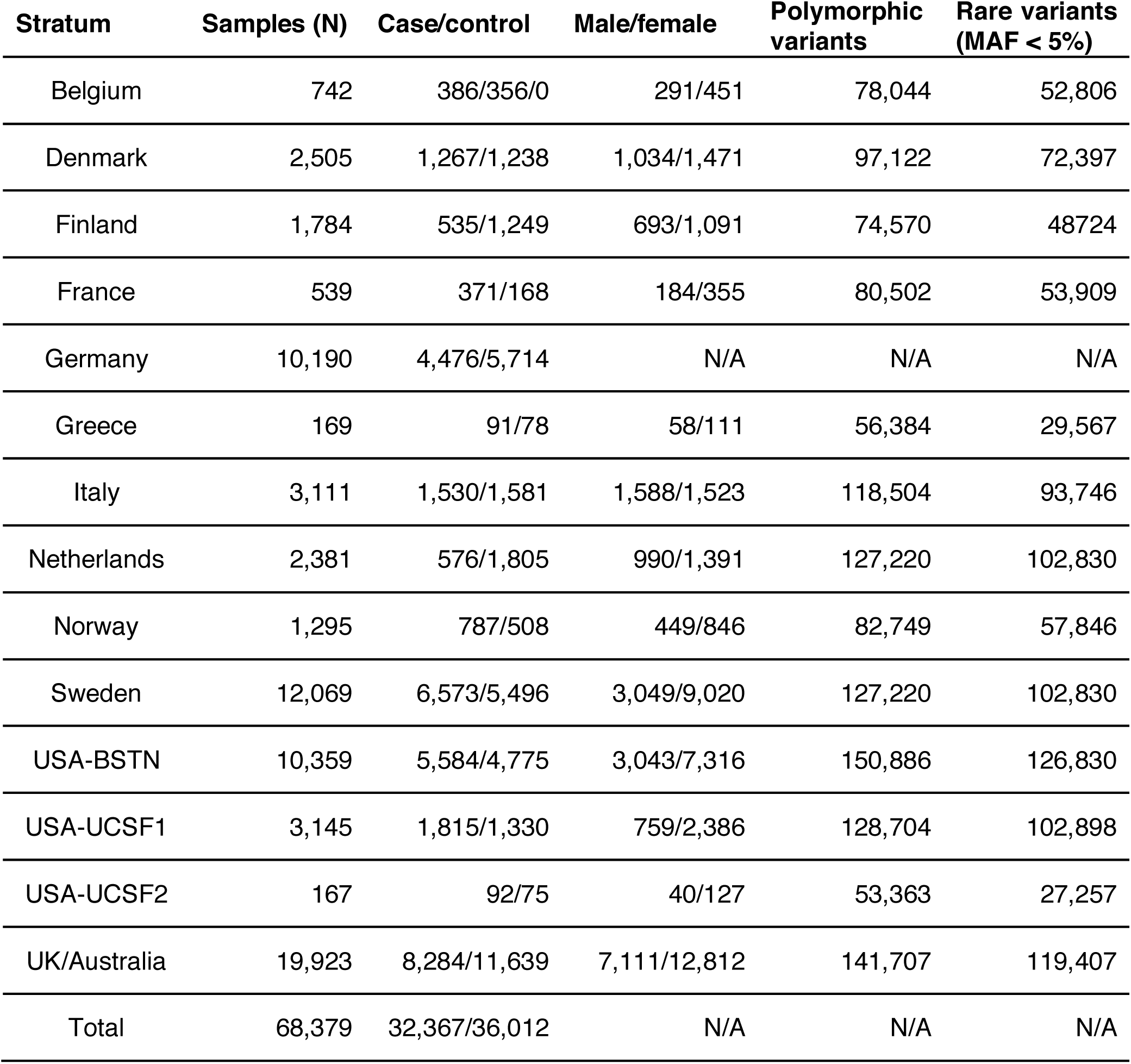
final stratum composition. We assigned 58,189 samples passing quality control to one of thirteen strata based on demography as described in the methods; data for 10,190 samples from Germany were received as post-QC summary statistics and are not included in Table S1. This gave us a total of 68,379 samples in our analysis. We had very few Australian control samples and so merged them with samples from the UK.

| Stratum | Samples (N) | Case/control | Male/female | Polymorphic variants | Rare variants (MAF < 5%) |
| --- | --- | --- | --- | --- | --- |
| Belgium | 742 | 386/356/0 | 291/451 | 78,044 | 52,806 |
| Denmark | 2,505 | 1,267/1,238 | 1,034/1,471 | 97,122 | 72,397 |
| Finland | 1,784 | 535/1,249 | 693/1,091 | 74,570 | 48,724 |
| France | 539 | 371/168 | 184/355 | 80,502 | 53,909 |
| Germany | 10,190 | 4,476/5,714 | N/A | N/A | N/A |
| Greece | 169 | 91/78 | 58/111 | 56,384 | 29,567 |
| Italy | 3,111 | 1,530/1,581 | 1,588/1,523 | 118,504 | 93,746 |
| Netherlands | 2,381 | 576/1,805 | 990/1,391 | 127,220 | 102,830 |
| Norway | 1,295 | 787/508 | 449/846 | 82,749 | 57,846 |
| Sweden | 12,069 | 6,573/5,496 | 3,049/9,020 | 127,220 | 102,830 |
| USA-BSTN | 10,359 | 5,584/4,775 | 3,043/7,316 | 150,886 | 126,830 |
| USA-UCSF1 | 3,145 | 1,815/1,330 | 759/2,386 | 128,704 | 102,898 |
| USA-UCSF2 | 167 | 92/75 | 40/127 | 53,363 | 27,257 |
| UK/Australia | 19,923 | 8,284/11,639 | 7,111/12,812 | 141,707 | 119,407 |
| Total | 68,379 | 32,367/36,012 | N/A | N/A | N/A |

## Acknowledgements

### Australia

Sample collection and genotyping was supported by NHMRC Project Grant APP605511.

### Belgium

BD, RL and PVD are Clinical Investigators of the Research Foundation Flanders (FWO-Vlaanderen). AG and BD are supported by the Research Fund KU Leuven (C24/16/045) and the Research Foundation Flanders (FWO-Vlaanderen) (G.0734.15). The SiGN study was funded by a cooperative agreement grant from the US National Institute of Neurological Disorders and Stroke, National Institutes of Health (U01-NS069208).

### Italy

Sample collection and genotyping was supported by the Italian Foundation for Multiple Sclerosis (FISM grants, Special Project “Immunochip” 2011/R/1, 2015/R/10) and Fondazione Cariplo (Grant 2010-0728).

### Norway

The Norwegian MS and control samples were collected and funded in collaboration between the Multiple Sclerosis research group in at Oslo University Hospital/University in Oslo, The Norwegian MS Registry and Biobank in Bergen and the Norwegian Bone Marrow Registry, supported by the Oslo MS Association, Bergen MS Society, Odda MS Society and the Research Council of Norway (grant 240102_Harbo).

### Sweden

Sample collection and genotyping was supported by the Swedish Medical Research Council; Swedish Research Council for Health, Working Life and Welfare, Knut and Alice Wallenberg Foundation, AFA insurance, Swedish Brain Foundation, the Swedish Association for Persons with Neurological Disabilities, Astra Zeneca Science for Life grant.

### UK

This study makes use of data generated as part of the Wellcome Trust Case Control Consortium 2 project (085475/B/08/Z and 085475/Z/08/Z), including data from the British 1958 Birth Cohort DNA collection (funded by the Medical Research Council grant G0000934 and the Wellcome Trust grant 068545/Z/02) and the UK National Blood Service controls (funded by the Wellcome Trust). The study was supported by the Cambridge NIHR Biomedical Research Centre, UK Medical Research Council (G1100125) and the UK MS society (861/07). We thank the National Institute for Health Research and NHS Blood and Transplant. TwinsUK is funded by the Wellcome Trust, Medical Research Council, European Union, the National Institute for Health Research (NIHR)-funded BioResource, Clinical Research Facility and Biomedical Research Centre based at Guy’s and St Thomas’ NHS Foundation Trust in partnership with King’s College London. We thank the volunteers from the Oxford Biobank(www.oxfordbiobank.org.uk) for their participation. The recall process was supported by the National Institute for Health Research (NIHR) Oxford Biomedical Research Centre Programme. We gratefully acknowledge the participation of all NIHR Cambridge BioResource volunteers, and thank the NIHR Cambridge BioResource centre and staff for their contribution. We thank the National Institute for Health Research and NHS Blood and Transplant.

### USA

We thank the Biorepository Facility and the Center for Genome Technology laboratory personnel (specifically Sandra West, Simone Clarke, Daniela Martinez, and Patrice Whitehead) within the John P. Hussman Institute for Human Genomics at the University of Miami for centralized DNA handling and genotyping for this project. Related funding support: The US National Multiple Sclerosis Society (grants RG-4680-A-1) and the NIH (R01-NS096212, R01-NS049477 and R01-NS026799).

## References

Altshuler, D., Daly, M.J., and Lander, E.S. (2008). Genetic Mapping in Human Disease. Science 322, 881–888.

Andlauer, T.F.M., Buck, D., Antony, G., Bayas, A., Bechmann, L., Berthele, A., Chan, A., Gasperi, C., Gold, R., Graetz, C., et al. (2016). Novel multiple sclerosis susceptibility loci implicated in epigenetic regulation. Sci Adv 2, e1501678–e1501678.

Australia and New Zealand Multiple Sclerosis Genetics Consortium (2009). Genome-wide association study identifies new multiple sclerosis susceptibility loci on chromosomes 12 and 20. Nature Genetics 41, 824–828.

Baranzini, S.E., Galwey, N.W., Wang, J., Khankhanian, P., Lindberg, R., Pelletier, D., Wu, W., Uitdehaag, B.M.J., Kappos, L., GeneMSA Consortium, et al. (2009). Pathway and network-based analysis of genome-wide association studies in multiple sclerosis. Hum. Mol. Genet. 18, 2078–2090.

Bettini, M.L., Pan, F., Bettini, M., Finkelstein, D., Rehg, J.E., Floess, S., Bell, B.D., Ziegler, S.F., Huehn, J., Pardoll, D.M., et al. (2012). Loss of Epigenetic Modification Driven by the Foxp3 Transcription Factor Leads to Regulatory T Cell Insufficiency. Immunity 36, 717–730.

Binder, M.D., Fox, A.D., Merlo, D., Johnson, L.J., Giuffrida, L., Calvert, S.E., Akkermann, R., Ma, G.Z.M., ANZgene, Perera, A.A., et al. (2016). Common and Low Frequency Variants in MERTK Are Independently Associated with Multiple Sclerosis Susceptibility with Discordant Association Dependent upon HLA-DRB1*15 : 01 Status. PLoS Genetics 12, e1005853.

Chami, N., Chen, M.-H., Slater, A.J., Eicher, J.D., Evangelou, E., Tajuddin, S.M., Love-Gregory, L., Kacprowski, T., Schick, U.M., Nomura, A., et al. (2016). Exome Genotyping Identifies Pleiotropic Variants Associated with Red Blood Cell Traits. Ajhg 99, 8–21.

Compston, A., and Coles, A. (2008). Multiple sclerosis. Lancet 372, 1502–1517.

Constant, S., Pfeiffer, C., Woodard, A., Pasqualini, T., and Bottomly, K. (1995). Extent of T cell receptor ligation can determine the functional differentiation of naive CD4+ T cells. J. Exp. Med. 182, 1591–1596.

Dankowski, T., Buck, D., Andlauer, T.F.M., Antony, G., Bayas, A., Bechmann, L., Berthele, A., Bettecken, T., Chan, A., Franke, A., et al. (2015). Successful Replication of GWAS Hits for Multiple Sclerosis in 10,000 Germans Using the Exome Array. Genet. Epidemiol. 39, 601–608.

de Jager, P.L., Jia, X., Wang, J., de Bakker, P.I.W., Ottoboni, L., Aggarwal, N.T., Piccio, L., Raychaudhuri, S., Tran, D., Aubin, C., et al. (2009). Meta-analysis of genome scans and replication identify CD6, IRF8 and TNFRSF1A as new multiple sclerosis susceptibility loci. Nature Genetics 41, 776–782.

Dominguez-Villar, M., Baecher-Allan, C.M., and Hafler, D.A. (2011). Identification of T helper type 1- like, Foxp3+ regulatory T cells in human autoimmune disease. Nature Medicine 17, 673–675.

Fuchsberger, C., Flannick, J., Teslovich, T.M., Mahajan, A., Agarwala, V., Gaulton, K.J., Ma, C., Fontanillas, P., Moutsianas, L., McCarthy, D.J., et al. (2016). The genetic architecture of type 2 diabetes. Nature 536, 41–47.

House, I.G., Thia, K., Brennan, A.J., Tothill, R., Dobrovic, A., Yeh, W.Z., Saffery, R., Chatterton, Z., Trapani, J.A., and Voskoboinik, I. (2015). Heterozygosity for the common perforin mutation, p.A91V, impairs the cytotoxicity of primary natural killer cells from healthy individuals. Immunol. Cell Biol. 93, 575–580.

Housley, W.J., Fernandez, S.D., Vera, K., Murikinati, S.R., Grutzendler, J., Cuerdon, N., Glick, L., De Jager, P.L., Mitrovic, M., Cotsapas, C., et al. (2015). Genetic variants associated with autoimmunity drive NFκB signaling and responses to inflammatory stimuli. Science Translational Medicine 7, 291–293.

International Multiple Sclerosis Genetics Consortium (2013). Analysis of immune-related loci identifies 48 new susceptibility variants for multiple sclerosis. Nature Genetics 45, 1353–1360.

International Multiple Sclerosis Genetics Consortium & Wellcome Trust Case Control Consortium 2 (2011). Genetic risk and a primary role for cell-mediated immune mechanisms in multiple sclerosis. Nature 476, 214–219.

Jakkula, E., Leppä, V., Sulonen, A.-M., Varilo, T., Kallio, S., Kemppinen, A., Purcell, S., Koivisto, K., Tienari, P., Sumelahti, M.-L., et al. (2010). Genome-wide association study in a high-risk isolate for multiple sclerosis reveals associated variants in STAT3 gene. Am. J. Hum. Genet. 86, 285–291.

Li, B., Samanta, A., Song, X., Iacono, K.T., Bembas, K., Tao, R., Basu, S., Riley, J.L., Hancock, W.W., Shen, Y., et al. (2007). FOXP3 interactions with histone acetyltransferase and class II histone deacetylases are required for repression. Proceedings of the National Academy of Sciences 104, 4571–4576.

Liu, D.J., Peloso, G.M., Yu, H., Butterworth, A.S., Wang, X., Mahajan, A., Saleheen, D., Emdin, C., Alam, D., Alves, A.C., et al. (2017). Exome-wide association study of plasma lipids in >300,000 individuals. Nature Genetics 49, 1758–1766.

Manolio, T.A., Collins, F.S., Cox, N.J., Goldstein, D.B., Hindorff, L.A., Hunter, D.J., McCarthy, M.I., Ramos, E.M., Cardon, L.R., Chakravarti, A., et al. (2009). Finding the missing heritability of complex diseases. Nature 461, 747–753.

Marouli, E., Graff, M., Medina-Gomez, C., Lo, K.S., Wood, A.R., Kjaer, T.R., Fine, R.S., Lu, Y., Schurmann, C., Highland, H.M., et al. (2017). Rare and low-frequency coding variants alter human adult height. Nature 542, 186–190.

Martinelli-Boneschi, F., Esposito, F., Brambilla, P., Lindström, E., Lavorgna, G., Stankovich, J., Rodegher, M., Capra, R., Ghezzi, A., Coniglio, G., et al. (2012). A genome-wide association study in progressive multiple sclerosis. Multiple Sclerosis Journal 18, 1384–1394.

Moutsianas, L., Jostins, L., Beecham, A.H., Dilthey, A.T., Xifara, D.K., Ban, M., Shah, T.S., Patsopoulos, N.A., Alfredsson, L., Anderson, C.A., et al. (2015). Class II HLA interactions modulate genetic risk for multiple sclerosis. Nature Genetics 47, 1107–1113.

Nelson, M.R., Wegmann, D., Ehm, M.G., Kessner, D., St Jean, P., Verzilli, C., Shen, J., Tang, Z., Bacanu, S.-A., Fraser, D., et al. (2012). An abundance of rare functional variants in 202 drug target genes sequenced in 14,002 people. Science 337, 100–104.

Nischwitz, S., Cepok, S., Kroner, A., Wolf, C., Knop, M., Müller-Sarnowski, F., Pfister, H., Roeske, D., Rieckmann, P., Hemmer, B., et al. (2010). Evidence for VAV2 and ZNF433 as susceptibility genes for multiple sclerosis. Journal of Neuroimmunology 227, 162–166.

Patsopoulos, N.A., the Bayer Pharma MS Genetics Working Group, the Steering Committees of Studies Evaluating IFNβ-1b and a CCR1-Antagonist, ANZgene Consortium, Gene MSA, International Multiple Sclerosis Genetics Consortium, and de Bakker, P.I.W. (2011). Genome-wide meta-analysis identifies novel multiple sclerosis susceptibility loci. Ann Neurol. 70, 897–912.

Patsopoulos, N., Baranzini, S.E., Santaniello, A., Shoostari, P., Cotsapas, C., Wong, G., Beecham, A.H., James, T., Replogle, J., Vlachos, I., et al. (2017). The Multiple Sclerosis Genomic Map: Role of peripheral immune cells and resident microglia in susceptibility. bioRxiv 143933.

Purcell, S.M., Moran, J.L., Fromer, M., Ruderfer, D., Solovieff, N., Roussos, P., O’Dushlaine, C., Chambert, K., Bergen, S.E., Kähler, A., et al. (2015). A polygenic burden of rare disruptive mutations in schizophrenia. Nature 506, 185–190.

Sadler, A.J., and Williams, B.R.G. (2008). Interferon-inducible antiviral effectors. Nat Rev Immunol 8, 559–568.

Sanna, S., Pitzalis, M., Zoledziewska, M., Zara, I., Sidore, C., Murru, R., Whalen, M.B., Busonero, F., Maschio, A., Costa, G., et al. (2010). Variants within the immunoregulatory CBLB gene are associated with multiple sclerosis. Nature Genetics 42, 495–497.

Sawcer, S., Ban, M., Maranian, M., Yeo, T.W., Compston, A., Kirby, A., Daly, M.J., de Jager, P.L., Walsh, E., Lander, E.S., et al. (2005). A high-density screen for linkage in multiple sclerosis. Ajhg 77, 454–467.

Sawcer, S., Franklin, R.J.M., and Ban, M. (2014). Multiple sclerosis genetics. The Lancet Neurology 13, 700–709.

Schoech, A., Jordan, D., Loh, P.-R., Gazal, S., O’Connor, L., Balick, D.J., Palamara, P.F., Finucane, H., Sunyaev, S.R., and Price, A.L. (2017). Quantification of frequency-dependent genetic architectures and action of negative selection in 25 UK Biobank traits. bioRxiv 188086.

Sims, R., van der Lee, S.J., Naj, A.C., Bellenguez, C., Badarinarayan, N., Jakobsdottir, J., Kunkle, B.W., Boland, A., Raybould, R., Bis, J.C., et al. (2017). Rare coding variants in PLCG2, ABI3, and TREM2 implicate microglial-mediated innate immunity in Alzheimer’s disease. Nature Genetics 49, 1373–1384.

The CHARGE Consortium Hematology Working Group (2016). Meta-analysis of rare and common exome chip variants identifies S1PR4 and other loci influencing blood cell traits. Nature Genetics 48, 867–876.

Wellcome Trust Case Control Consortium (2007). Association scan of 14,500 nonsynonymous SNPs in four diseases identifies autoimmunity variants. Nature Genetics 39, 1329–1337.

Westerlind, H., Ramanujam, R., Uvehag, D., Kuja-Halkola, R., Boman, M., Bottai, M., Lichtenstein, P., and Hillert, J. (2014). Modest familial risks for multiple sclerosis: a registry-based study of the population of Sweden. Brain 137, 770–778.

Yang, J., Lee, S.H., Goddard, M.E., and Visscher, P.M. (2011). GCTA: A Tool for Genome-wide Complex Trait Analysis. The American Journal of Human Genetics 88, 76–82.

Zeng, J., de Vlaming, R., Wu, Y., Robinson, M., Lloyd-Jones, L., Yengo, L., Yap, C., Xue, A., Sidorenko, J., McRae, A., et al. (2017). Widespread signatures of negative selection in the genetic architecture of human complex traits. bioRxiv 145755.

## Methods references

Abraham, G., and Inouye, M. (2014). Fast Principal Component Analysis of Large-Scale Genome-Wide Data. PloS One 9, e93766.

Goldstein, J.I., Crenshaw, A., Carey, J., Grant, G.B., Maguire, J., Fromer, M., O’Dushlaine, C., Moran, J.L., Chambert, K., Stevens, C., et al. (2012). zCall: a rare variant caller for array-based genotyping: Genetics and population analysis. Bioinformatics 28, 2543–2545.

Lek, M., Karczewski, K.J., Minikel, E.V., Samocha, K.E., Banks, E., Fennell, T., O’Donnell-Luria, A.H., Ware, J.S., Hill, A.J., Cummings, B.B., et al. (2016). Analysis of protein-coding genetic variation in 60,706 humans. Nature 536, 285–291.

Price, A.L., Patterson, N.J., Plenge, R.M., Weinblatt, M.E., Shadick, N.A., and Reich, D. (2006). Principal components analysis corrects for stratification in genome-wide association studies. Nature Genetics 38, 904–909.

Purcell, S., Neale, B., Todd-Brown, K., Thomas, L., Ferreira, M.A.R., Bender, D., Maller, J., Sklar, P., de Bakker, P.I.W., Daly, M.J., et al. (2007). PLINK: a tool set for whole-genome association and population-based linkage analyses. Ajhg 81, 559–575.

